# Optimizing the design of spatial genomic studies

**DOI:** 10.1101/2023.01.29.526115

**Authors:** Andrew Jones, Diana Cai, Didong Li, Barbara E. Engelhardt

## Abstract

Spatially-resolved genomic technologies have shown promise for studying the relationship between the structural arrangement of cells and their functional behavior. While numerous sequencing and imaging platforms exist for performing spatial transcriptomics and spatial proteomics profiling, these experiments remain expensive and labor-intensive. Thus, when performing spatial genomics experiments using multiple tissue slices, there is a need to select the tissue cross sections that will be maximally informative for the purposes of the experiment. In this work, we formalize the problem of experimental design for spatial genomics experiments, which we generalize into a problem class that we call *structured batch experimental design*. We propose approaches for optimizing these designs in two types of spatial genomics studies: one in which the goal is to construct a spatially-resolved genomic atlas of a tissue and another in which the goal is to localize a region of interest in a tissue, such as a tumor. We demonstrate the utility of these optimal designs, where each slice is a two-dimensional plane, on several spatial genomics datasets.

## 1 Introduction

Spatially-resolved genomic assays present an opportunity to study the physical organization of cells, and how cell phenotype varies across space (Moses and Pachter, 2022). These assays have been used to study a variety of biological tissues and organs, such as brain (Lein et al., 2007; Zhang et al., 2021), liver (Saviano et al., 2020), heart (Mantri et al., 2021), and various tumors (Smith and Hodges, 2019).

However, spatial genomics experiments are costly in terms of financial, labor, and material resources. Ideally, an experiment studying a particular tissue type would collect genomic data— gene expression, protein expression, etc.—from the entire spatial domain of the tissue of interest. However, cost-constrained scientists are typically forced to select only a small fraction of the tissue to profile—the *field of view*. A common approach is to identify an anatomical region within the tissue of interest and take one or multiple parallel cross-section slices of the tissue in the region of interest. This approach is attractive for its simplicity in collecting the slices and for its adherence to orthodox slicing strategies (i.e., slicing along primary anatomical axes, such as coronal or sagittal axes).

Despite its attractive properties, this data collection approach may not provide the most informative data. In particular, adjacent, parallel cross-sections may contain largely redundant information that could be imputed from more distant, potentially non-parallel slices. Thus, there is a need for a systematic approach to designing spatial genomics experiments—in particular, optimizing the choice of which tissue cross-sections to collect—such that the experiments are maximally informative within the experimenter’s cost constraints.

Here, we propose a statistical approach to optimizing experimental design for spatial genomics studies. Specifically, we focus on the problem of choosing which cross sections of a tissue will yield a maximally informative experiment when these sections are profiled with a spatial genomics assay. Our proposed approach relies on fundamental concepts in Bayesian optimal experimental design (BOED) (Chaloner and Verdinelli, 1995). For a given statistical model of the data, our method finds the slice that is expected to provide the maximum amount of additional information about the tissue of interest. Our framework allows for designing experiments with a variety of experimental goals and can be adapted to the experimenter’s preferred statistical modeling approach. We demonstrate our approach through two different applications: building a tissue atlas and localizing a tissue region containing a tumor.

### 1.1 Related work

#### 1.1.1 Optimal experimental design

The literature on optimizing and automating experimental design has a long history (Russell, 1926; Fisher, 1926; Lindley, 1956; Box, 1980).

Optimal design first arose in the frequentist literature, and specifically in the setting of linear models (see (Steinberg and Hunter, 1984) for an early review). Frequentist approaches to experimental design start by positing an optimality criterion, which is defined by a loss functional. Let **X** be an *n* × *p* design matrix where the experimenter is tasked with choosing *n* designs from a design space 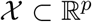. A design **X**\* is optimal with respect to a criterion with loss functional 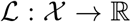 if it satisfies 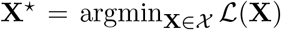. Several criteria have been proposed, such as *D*-optimality, where 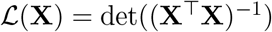; *A*-optimality, where 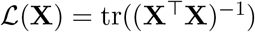; and *E*-optimality, which maximizes the top eigenvalue of (**X**^⊤^**X**)^-1^.

Bayesian optimal experimental design (BOED) (Chaloner and Verdinelli, 1995) extends experimental design to the setting of Bayesian inference, where the model parameter *θ* is assumed to be random and drawn from a prior distribution *π*(*θ*), and each observation *y* is drawn from a likelihood model *p*(*y*|*θ,x*). Given a vector of n observations 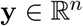, standard Bayesian inference proceeds by computing the posterior distribution, *p*(*θ*|y). Similar to the frequentist approach to experimental design, BOED techniques seek designs **X** that are expected to improve statistical inference in some way. However, unlike the frequentist setting that optimize a functional of a frequentist estimator, BOED approaches seek to improve the posterior according to some criterion. One of the most popular criteria is the expected information gain (EIG), which is defined as the expected difference in entropy between the prior and posterior:

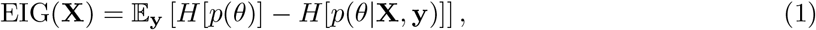

where *H*[*p*(*ω*)] = — *∫_Ω_p*(*ω*)log*p*(*ω*)*dω* is the differential entropy of a density function *p*(*ω*). A design that maximizes the EIG is also sometimes called Bayesian *D*-optimal because, in the case of a linear model, maximizing the EIG is equivalent to finding a *D*-optimal design. The EIG can also be written in terms of predictive distributions in place of posterior distributions (see Appendix A.1.1 for a derivation):

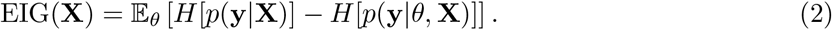

Depending on the setting, one form of the EIG can be easier to work with than the other. For complex statistical models, computing the posterior and predictive distributions analytically is impossible, therefore making the EIG computationally intractable as well. Recently, approximate inference methods for BOED have been proposed that ease the computational burden (Foster et al., 2019, 2020, 2021). However, these approaches come at the expense of an exact solution.

#### 1.1.2 Experimental design for genomics studies

In genomics, automating experimental design has become of special interest in recent years due to the rising cost and complexity of experimental protocols. Several statistical approaches have been proposed for designing single-cell sequencing experiments (Svensson et al., 2017; Camerlenghi et al., 2020; Schmid et al., 2020; Masoero et al., 2022).

In the field of spatial genomics, a technique was recently developed for determining the appropriate experimental parameters to achieve a desirable level of statistical power (Baker et al., 2022). Despite these advancements, there remains a lack of methods for optimizing the physical locations of a tissue to profile.

## 2 Notation and problem statement

We now formalize the problem studied in this paper. We consider a spatial genomics dataset consisting of pairs (**x**, **y**), where 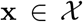 is a spatial location, 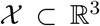 is the spatial domain of the tissue (we are assuming it is three dimensional), and **y** is a *p*-dimensional outcome at this location (e.g., a vector of gene expression, protein expression, or another univariate or multivariate phenotype at this location). We denote a dataset of *n* such pairs as 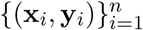. In matrix form, we define **X**_*t*_ and **Y**_*t*_ to be the *n* × 3 and *n*×*p* matrices whose ith rows are the **x**_*i*_ and **y**_*i*_, respectively.

Suppose the goal of an experiment is to profile the phenotype of a tissue, organ, or entire organism whose cells lie in the spatial domain 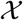. Spatially-resolved genomic data is typically collected from *slices* of the tissue. Each slice is a two-dimensional cross-section of 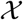. Let *P* represent a plane intersecting the tissue; the specific parameterization for the plane is not critical. The set of points on a cross section defined by *P* is given by 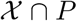. In practice, tissue sections have nonzero width, meaning that points nearby a cross section’s plane will also be observed. For a slice with half-width *δ*, we denote the set of points observed by a cross section defined by plane *P* as 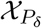; these are the spatial locations within a Euclidean distance of *δ* to the plane *P*.

We consider the class of iterative experimental strategies, where spatial genomics readouts are collected in *T* sequential batches, and each batch is a single experiment that collects data from one tissue slice. When planning batch *t* ∈ [*T*], our goal is to select a cross section defined by *P_t_* that maximizes the expected utility gained from performing that experiment. Let 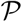 denote the set of candidate planes, and let *θ* ∈ Θ be a set of unknown model parameters. Throughout this paper, 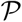 will constitute our *design space*, or the set of designs to be chosen from. We define 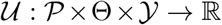 to be a utility function that, intuitively, measures the goodness of an experiment and its expected observations. We discuss the choice of the utility function in the next section. On the first batch, the optimal slice is given by

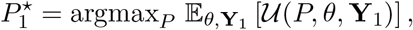

where **Y**_1_ is the set of outcomes observed on the first iteration. The maximizer for batches *t* > 1 is similar but also relies on the data collected up to that point to inform the design of the current batch. For *t* > 1 the solution is

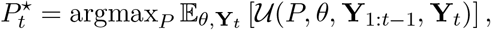

where **Y**_*t*_ are the outcomes observed on iteration *t* and **Y**_1:*t*_ = {**Y**_1_,…, **Y**_*t*_}.

An important aspect of this design problem—and one that makes it unique from related experimental design problems—is its highly structured design space. Specifically, while other design problems allow the experimentalist to freely choose one or multiple designs (unique values of x) on each iteration, the spatial slicing problem requires that the spatial locations be situated on the same plane. We refer to this general problem class as *structured batch experimental design*, which encompasses experimental settings where multiple samples are collected in a constrained manner on each iteration. In our case, the samples are constrained to lie on a plane. Other fields in which structured batch experimental design might appear are tomography (Buzug, 2011) and pathology (Kierszenbaum and Tres, 2015).

## 3 Methods

We now describe our approach to the spatial experimental design problem. We consider two possible experimental goals:

1. Building a spatially-resolved genomic atlas for a tissue or organ;
2. Localizing a tissue region of interest, such a tumor or anatomical region.

For each of these applications, we formalize the experimental goal, and we propose a statistical model and utility function that reflect the associated goal. We then propose an optimization scheme to iteratively find the sample cross section with maximum expected utility.

### 3.1 Atlas building objective

A long-term goal in genomics and biology is to build a comprehensive characterization of all cell types in the human body. This is commonly referred to as an *atlas*, which draws an analogy with a “map” of cells’ physical organization and phenotypes (Quake, 2022). In its ideal form, a comprehensive atlas would allow researchers to query the atlas using a spatial location or region of interest, and the query would return a detailed description of the phenotype, including cell types and cell states.

Atlases for a set of human and mouse tissue types have been constructed using various data modalities, such as *in situ* hybridization, single-cell RNA-sequencing, histology, and spatial gene expression (Lein et al., 2007; Sunkin et al., 2012; Rozenblatt-Rosen et al., 2017; Quake, 2022; Shekhar and Sanes, 2021). However, comprehensive atlases using the most modern spatial gene expression profiling methods have yet to be established.

Here, we consider the problem of efficiently constructing an atlas using spatial genomics technologies. We first formalize the problem of building a spatially-resolved atlas and then discuss our proposed approach. For simplicity, we first consider a noisy univariate phenotype *y* (e.g., the expression level of one gene) and move to multivariate phenotypes later. Suppose *y* follows a spatial process defined on the domain 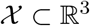. Consider the following model:

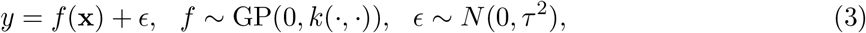

where *f* has a Gaussian process prior with mean zero and covariance function *k*(**·, ·**), and *ϵ* is Gaussian noise with variance *τ*^2^. Under this model, the unknown function *f*(**·**) provides a full description of the spatially-resolved atlas for phenotype *y*. In particular, given spatial location **x**, the function evaluation *f* (**x**) tells us the (noiseless) value of *y* at that location. Thus, the statistical goal in atlas-building is to infer *f* (**x**) for every location **x** within the 3D domain. We take a Bayesian approach to this problem, where estimating the function *f* (**·**) amounts to computing the posterior distribution for *f* given the collected data,

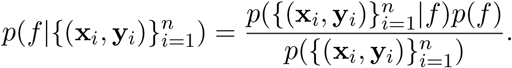

Our experimental design objective is then to choose tissue slices that are expected to maximally “improve” this posterior distribution in some way. We discuss metrics to quantify this improvement next.

#### 3.1.1 Atlas construction via information gain

Since our ultimate goal in the atlas-building objective is to infer the unobserved function *f*, we choose a utility function that rewards experimental designs that offer more information about *f*. The *information gain* (IG) is a utility function defined as the difference in entropy between the prior and the posterior distributions. Under our atlas model, the IG of taking a slice defined by plane *P_t_* and observing outcome **Y**_*t*_ is IG(**Y**_*t*_, *P_t_*) = *H*[*p*(*f* |**Y**_1:*t*-1_)] — *H*[*p*(*f* |**Y**_1:*t*_,*P_t_*)], where *H*[**·**] is the differential entropy functional. Because **Y**_*t*_ is not observed before conducting experiment *t*, we cannot directly optimize the IG with respect to *P_t_*. Thus, we take the expectation of the IG with respect to **Y**_*t*_, which is a quantity known as the *expected information gain* (EIG). The EIG of design *P_t_* is given by

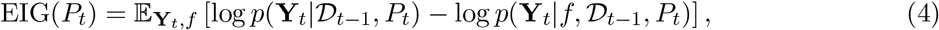

where 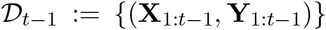 represents the data observed through experimental iterations 1,…, 1 — *t*, and **X**_1:*t*_ and **Y**_1:*t*_ are the spatial locations and associated outcomes, respectively. We then maximize the EIG (Equation 4) with respect to *P_t_*.

Under our GP regression atlas model (Equation 3), the EIG can be computed analytically. Let **X**_*t*_ be the set of spatial locations captured on the cross section defined by plane *P_t_*. The EIG for *P_t_* is

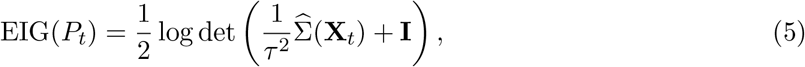

where the predictive covariance 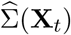 of the GP at locations **X**_*t*_ is given by

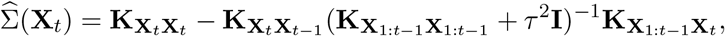

and we use the notation **K_XX’_** to denote the matrix of covariance function evaluations whose *ij*th element is 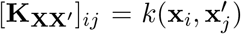. See Appendix A.1.2 for a more detailed derivation of these quantities. Our optimization problem under the atlas-building objective is to maximize the EIG with respect to *P_t_*.

#### 3.1.2 Maximizing information gain to find the optimal cross section

Our goal is to find the set of points in a tissue that maximize the EIG (Equation 5). However, because we are constrained to collect two-dimensional cross sections of a tissue, rather than any arbitrary subset of spatial locations, we must constrain each candidate design’s spatial locations **X** to lie on a plane.

This is equivalent to choosing a plane *P_t_* representing a two-dimensional cross-section of the slice collected at time *t*. Although there are an infinite number of potential cross sections, we simplify the optimization problem by discretizing the space of cross sections. Specifically, we create a design space with *D* cross sections, where *D* can be chosen depending on computational resources and required precision. The optimization problem then reduces to maximizing over a discrete set, which in this setting is typically a tractable problem.

Because slices consist of entire cross sections of the tissue, each slice creates a new, disjoint tissue fragment. At the start of iteration *t*, we will have taken *t* — 1 slices, so the tissue will be split into *t* disjoint fragments. We account for this by considering the candidate cross sections in each fragment separately. Let 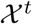 be the set of spatial locations corresponding to tissue fragment *t*, and let 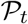 be the set of planes intersecting 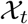. Note that, by definition, for all *t* it must hold that

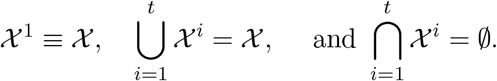

The optimization problem on iteration t is then to find the slice that maximizes the EIG for the current set of tissue fragments: 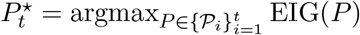.

### 3.2 Localizing a tissue region of interest

Next, we consider an experimental design setting in which there is a particular region of space within 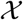 whose borders we would like to identify. For example, we may be studying a biopsy sample from a tumor tissue, and we would like to identify the border between the tumor and healthy tissue using as few slices as possible. This localization problem is a common goal in cancer pathology (Yoosuf et al., 2020).

For simplicity, assume that we can label each spatial location as being on the interior of the region of interest (*y* = 1) or the exterior of the region of interest (*y* = 0) after we have collected data for that location. On experimental iteration *t*, we collect data from a cross section of the tissue defined by *P_t_*, where the data are made up of the spatial locations 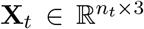 and the interior/exterior labels for those locations **y**_*t*_ ∈ {0,1}^*nt*^, where *n_t_* is the number of points observed after taking slice *P_t_*.

**Bounding box model** Consider a logistic regression model where the area of interest is modeled with an axis-aligned rectangular bounding box. Assume a spatially-varying Bernoulli likelihood, *y* ~ Bern(*g*(**x**)), where *g*(**·**) is a link function mapping the spatial coordinates to the bounding box probability model. For the axis-aligned bounding box, we can parameterize *g*(**·**) as follows:

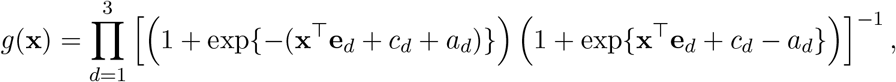

where 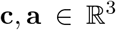 are parameters controlling the center and width of the box, respectively, and **e**_*d*_ is the dth axis-aligned unit vector of length 3. Isotropic Gaussian priors can be used, i.e., **a**, **c** ~ *N*(**0**,**I**). This model, which is a generalization of a logistic regression model, captures the borders of the region of interest through the parameters *θ* = {**c**, **a**}. Thus, the posterior after iteration *t* is *p*(*θ*|**X**_1:*t*_,**y**_1:*t*_). Recall that 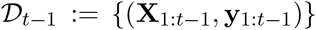 represents the data observed through experimental iterations 1,…, *t* — 1. The expected information gain for a slice through plane *P_t_* is

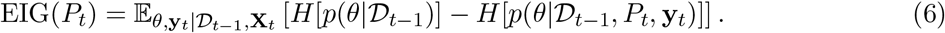

In order to select the slices to identify the borders of the region of interest representing the tumor, we maximize the EIG with respect to P_t_.

**Spherical and elliptical border model** We can parameterize the border of a region of interest using shapes other than a rectangle. For example, we can use a circular bounding area instead. Recall that the equation for the points in 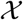 contained within a ball with center **c** and radius *r* is given by 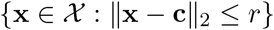. A viable statistical model is then

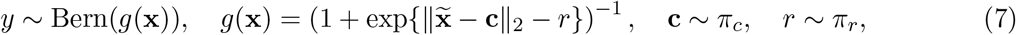

where *π_c_* and *π_r_* are prior distributions for the center and radius, respectively.

The spherical border model can also be generalized to an elliptical border. Recall that an ellipsoid can be written as a linear transformation of a sphere. The points contained inside the ellipsoid are 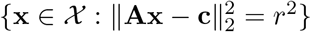, where 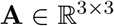. These options for border-finding extend our experimental design approach.

#### 3.2.1 Inference

Under any of these border-finding models, our goal is to find the design that maximizes the EIG (Equation 4). For most modeling settings, there are three intractable quantities in the expression for EIG: the posterior 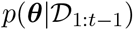, the entropy of the posterior, and the outer expectation over the data.

We use a two-step approximation for these intractable quantities. First, while designing iteration *t*, we compute an approximation to the posterior 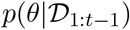 using variational inference. We denote this approximation as *q*_*t*-1_(*θ*) (see Appendix A.1.3 for details). Second, we use this approximate posterior to estimate the EIG using a nested Monte Carlo (NMC) sampling approach (Rainforth et al., 2018).

We briefly review the NMC estimator to solve this problem. Expanding the EIG expression, we see that it contains a nested integral:

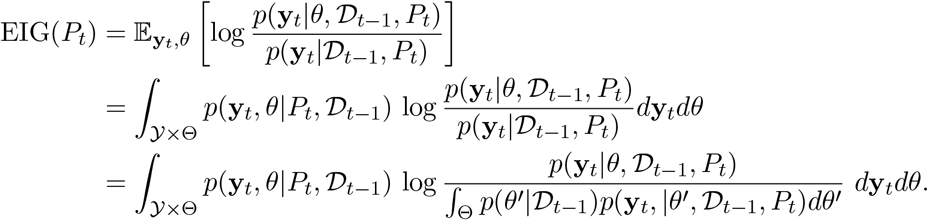

The NMC estimator (Rainforth et al., 2018) approximates these nested integrals using a Monte Carlo estimator. Taking *S* samples for the outer sum and *S’* samples for the inner sum, the NMC approximation is

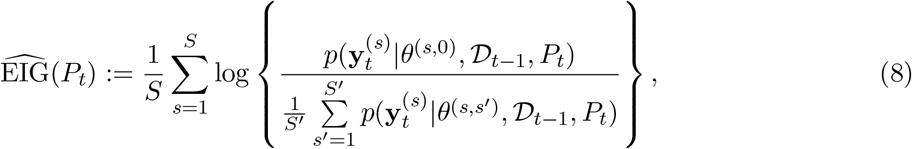

where *θ*^(*s*,0)^ and *θ*^(*s,s’*)^ are sampled from a variational approximation to the posterior for *θ*, and 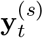 is sampled from the likelihood model:

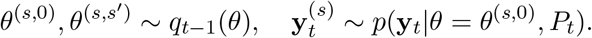

Here, *q*_*t*-1_(*θ*) is a variational approximation to the posterior 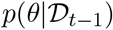 (see Appendix A.1.3 for details). Algorithm 1 contains a more detailed exposition of the approach. Increasing the number of Monte Carlo samples *S*, *S’* trades off computation speed for a more precise estimate of the EIG. Selecting the optimal slice is then performed via 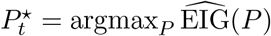.

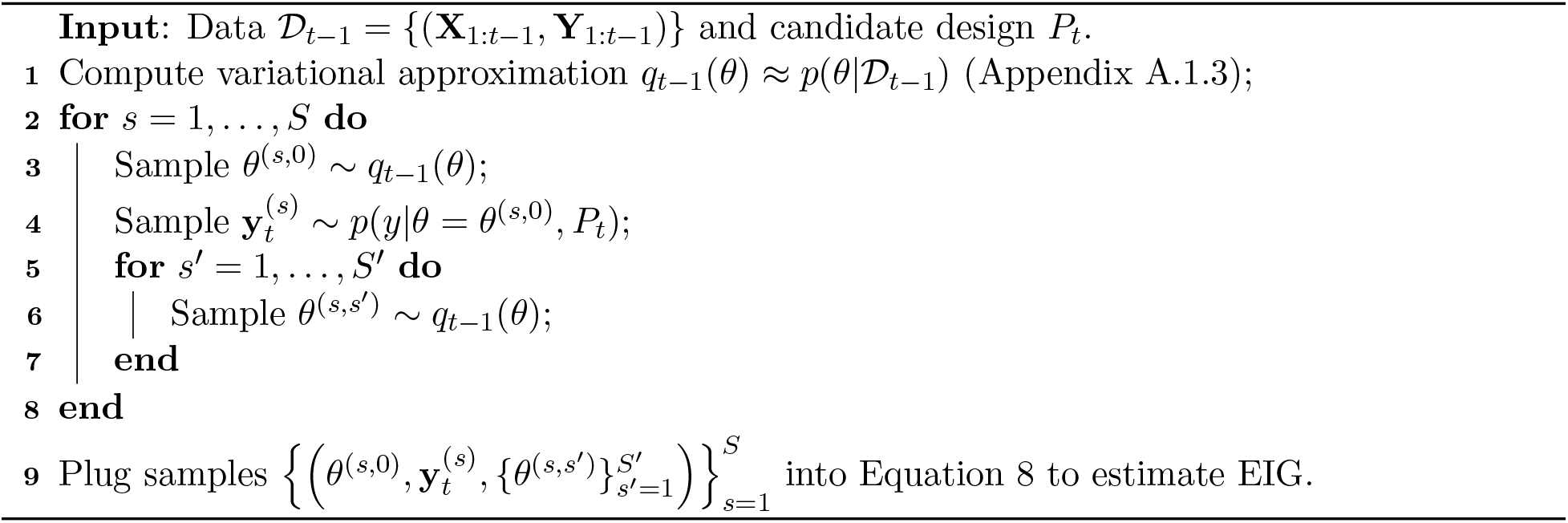

## 4 Experiments

### 4.1 Experimental setup

We now demonstrate our experimental design approach through applications to synthetic data and three spatial gene expression datasets. Throughout our experiments, we compare five methods for experimental design:

- *EIG*: Maximize EIG over candidate cross sections while accounting for tissue fragmenting.
- *EIG (parallel)*: Maximize EIG over candidate cross sections while constraining the slices to be parallel and along an anatomical axis.
- *EIG (no fragmenting)*: Maximize EIG over candidate cross sections, allowing for slices to cut across multiple tissue fragments.
- *Serial*: Take serial parallel cross sections along an anatomical plane.
- *Random*: Randomly choose from candidate cross sections while accounting for tissue fragmenting.

The first three approaches, *EIG, EIG (parallel)*, and *EIG (no fragmenting)* are special cases of our proposed approach with different design spaces. The *Serial* design approach is the one most commonly used in spatial genomics experiments. The *Random* approach would never be used in practice but serves as a baseline comparison.

In the following experiments, we consider cross sections through a tissue, where the tissue is represented by a cloud of points. As described above, each cross section is defined by a plane. We define a slice’s width as *δ* and allow any spatial location within distance *δ* to the plane to be observed after taking this slice. In practice, the choice of the section width *δ* is often dictated by the data collection modality being used, as well as the tools available.

### 4.2 Simulations

We first conducted a series of simulation studies in order to evaluate the behavior of our approach to structured batch experimental design.

#### 4.2.1 Small-scale demonstration

As an initial demonstration and visualization of our approach, we applied our method for atlas building to a simulated tissue. For ease of visualization, we first study a setting where we are tasked with choosing one-dimensional slices (lines) from a two-dimensional tissue. The simulated tissue had a circular shape with a radius of five (Figure 1a). We placed points randomly within the boundaries of the tissue, which represent the locations of cells or spots. We then generated synthetic responses at each location using a GP with a mean of zero and Matérn 1/2 covariance function with lengthscale *ℓ* = 1 and noise variance *τ*^2^ = 0.1.

**Figure 1:**
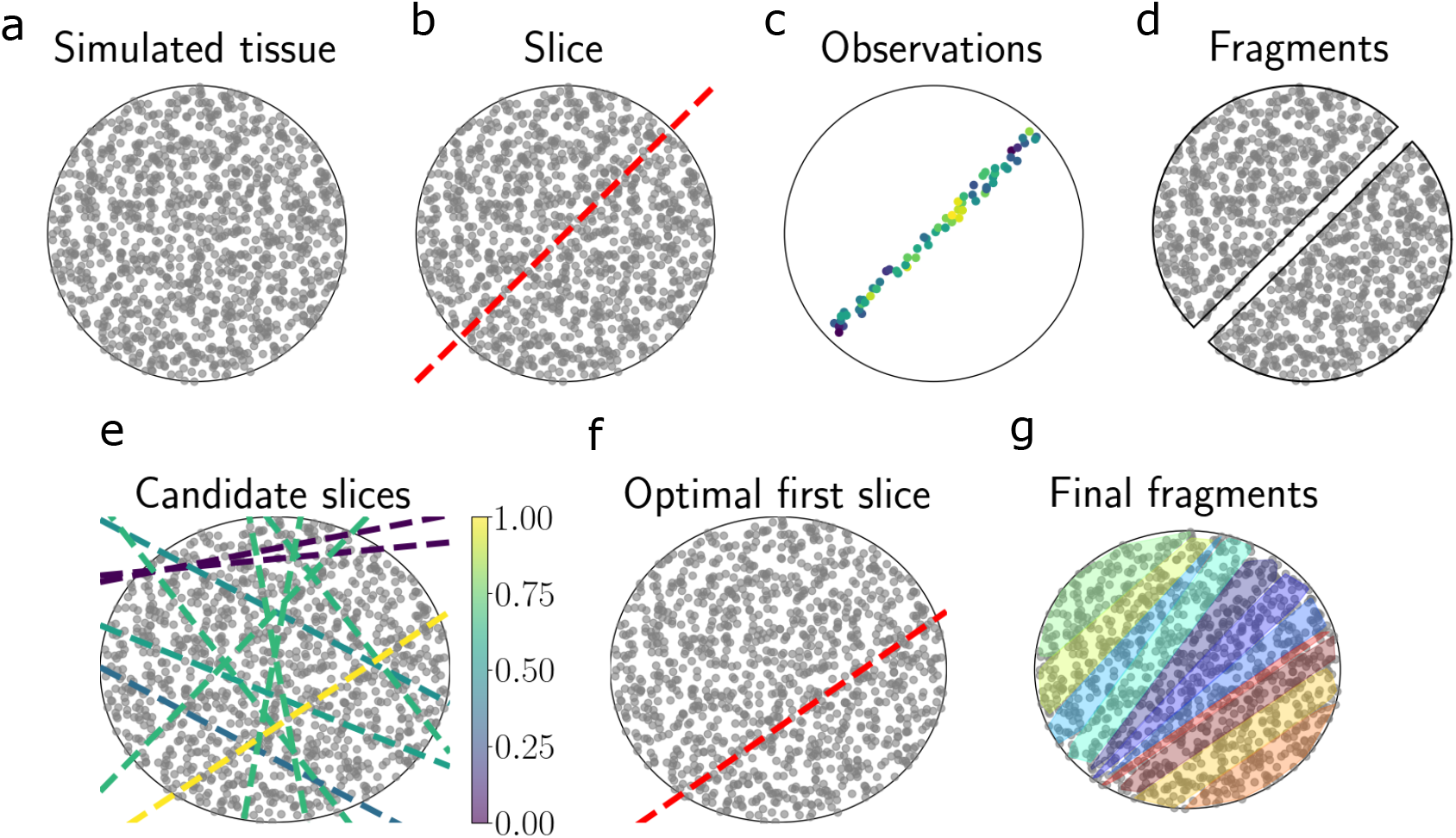
Demonstration of slicing in two-dimensional simulated tissue. (a) Simulated spherical tissue with a grid of spots. (b) An example one-dimensional slice through the tissue. (c) The resulting observations at each spot after taking the slice in (b). The colors represent a univariate phenotype. (d) After slicing, the simulated tissue is split into two fragments. (e) Each line represents a candidate one-dimensional slice. Each slice is colored by its EIG (normalized to have a maximum of one). (f) The EIG-maximizing slice. (g) Tissue fragments after *T* = 10 iterations of slicing. Each color represents a distinct fragment.

On each iteration of this experiment, the objective was to choose a one-dimensional cross-section through the synthetic tissue. After taking a slice, the observations from cells lying within distance *δ* of the slice’s line, along with the response value at each of these cells, were revealed. The tissue then broke into two *T* + 1 fragments (Figure 1b-d). We then repeated this process for a total of *T* iterations.

We applied our experimental design approach to this problem to find the EIG-optimal experimental design on each iteration. To do so, we first discretized the space of possible designs. We parameterized each design by its angle *φ* with the *x*-axis and its intercept *b*_0_ with the *y*-axis, and took 50 slopes whose angles are equally spaced in [0,*π*) and 50 intercepts equally spaced in [−5, 5]. (The slope of the line is given by tan *φ*.) Using all pairwise combinations of *φ* and *b*_0_, this resulted in a design space containing 2500 cross sections.

We ran our method forward for *T* = 10 iterations. On each iteration, we computed the EIG for each possible slice and selected the slice with the highest EIG (Figure 1e). We then visualized the resulting slices.

We found that, in general, the optimal slices under our criterion tended to be the cross sections that intersected the most cells (Figure 1g). On the first iteration, a slice near the center of the circular domain was chosen. On subsequent iterations, slices that were somewhat parallel to the first slice were chosen. This demonstration suggests that EIG maximization under the atlas model encourages choosing slices including many cells over slices containing few cells.

#### 4.2.2 Three-dimensional demonstration

We next conducted a similar experiment, but this time we extended it to a three-dimensional spatial domain. We placed points randomly within a cube with edge length 10 and generated synthetic responses at each location from a GP with a radial basis function (RBF) covariance function with length scale *ℓ* = 1 and noise variance *τ*^2^ = 0.1 (Figure 2a). We ran our experimental design procedure for *T* = 10 iterations. To quantitatively evaluate the chosen cross sections, we ran a prediction experiment on each iteration. Specifically, after iteration *t*, we fit a GP with an RBF covariance function using the data collected theretofore, estimating the RBF hyperparameters using maximum likelihood estimation. We then computed the predictive mean for the unobserved spots and computed the goodness-of-fit *R*^2^ between the predictions and the true values. Intuitively, we expect a better design procedure to select slices that will yield better predictive ability, therefore allowing more efficient imputation of the “atlas”. We compared our *EIG* experimental design method to the *Random* approach and the *Serial* approach.

**Figure 2:**
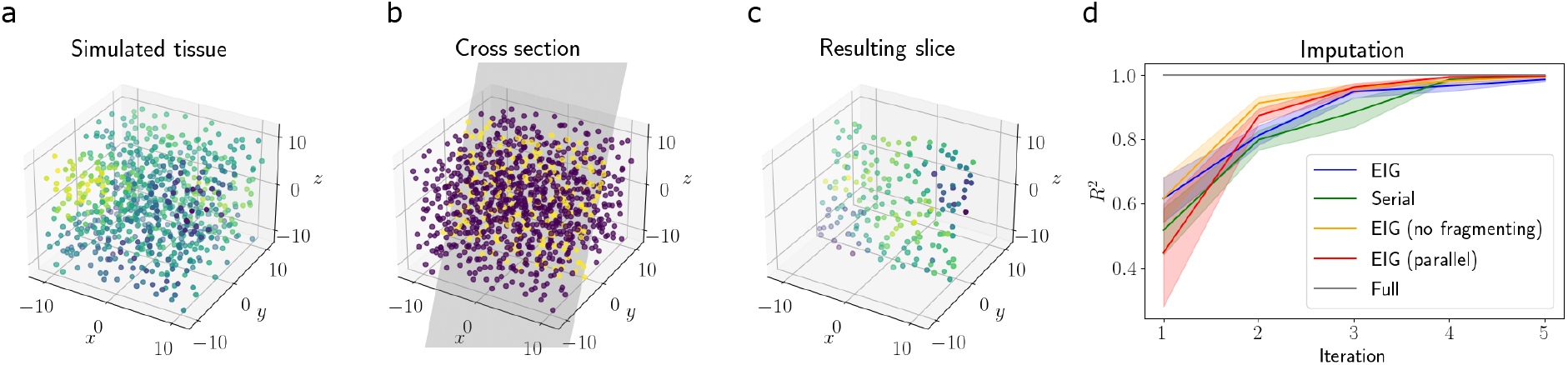
Imputing unobserved gene expression from observed cross sections. (a) Simulated tissue colored by synthetic gene expression. (b) An example slice through the synthetic tissue. (c) The resulting observations from the slice in (b). (d) *R*^2^ for imputations of gene expression after each slicing iteration for each method.

We found that our design procedure yielded improved predictive performance compared to the competing approaches (Figure 2d). Specifically, the *R*^2^ of the predictions for the *EIG* design method approached the *R*^2^ level of the complete atlas in fewer experimental iterations than competing approaches. The *EIG* method reached the performance of the full atlas after collecting roughly four slices, while the *Serial* method required roughly six slices to achieve comparable performance. This result suggests that the *EIG* approach selects cross sections that allow for efficient construction of cell atlases.

#### 4.2.3 Border finding

We next studied a simulated setting in which the goal is to identify the location and boundaries of a tissue region of interest. This simulation mimics several applications in spatial genomics, such as identifying the boundaries of a tumor and localizing an anatomical region of interest. We generated a dataset with two-dimensional spatial coordinates, similar to the data generation in Section 4.2.1. On the interior of the synthetic tissue, we designated points within a circle as the region of interest (ROI) (Figure 3a). The points outside of this circle were given a label of region of non-interest. We injected noise to these labels so that 10% of all points were mislabeled. The goal in this experiment was to collect cross sections of points in order to localize the ROI as quickly as possible.

**Figure 3:**
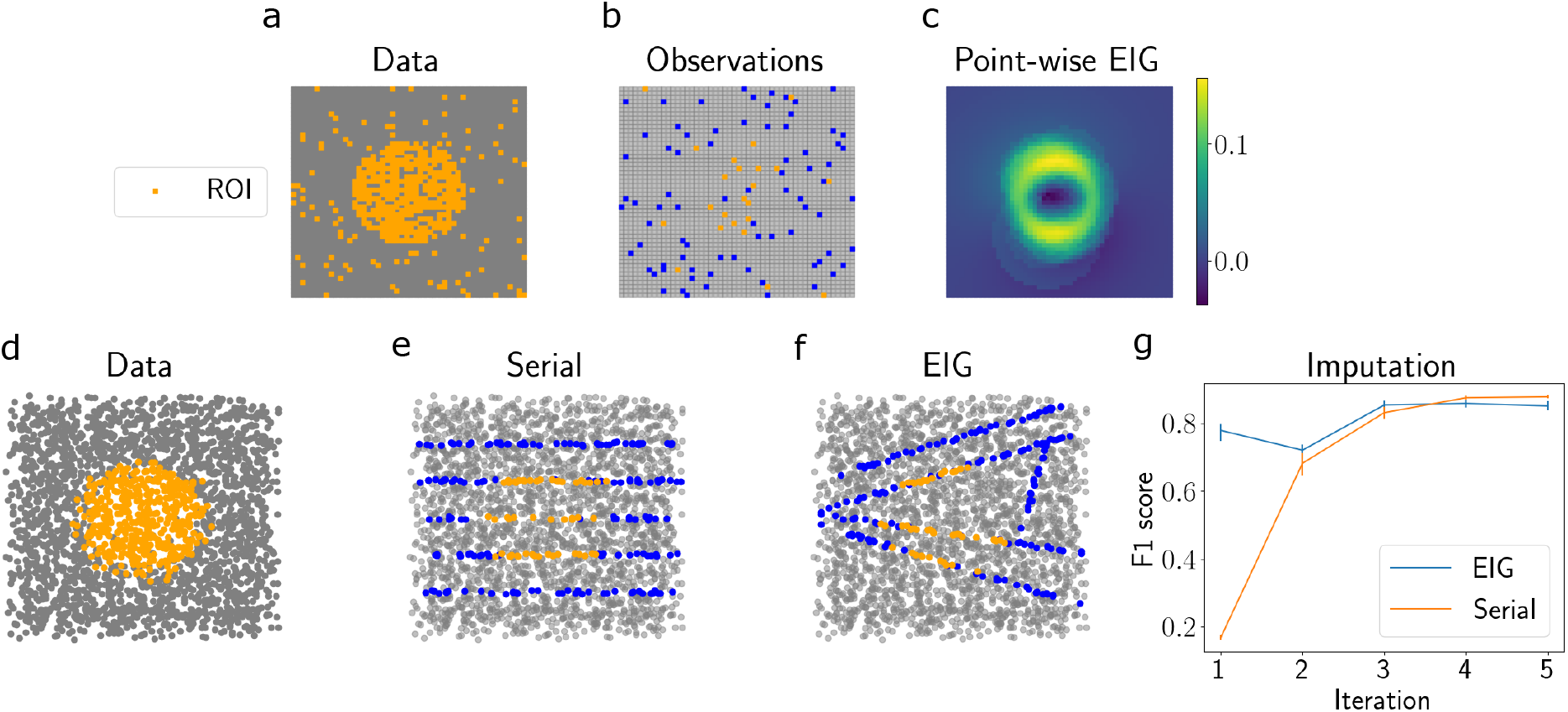
Synthetic slicing experiment for localizing a region of interest. (a) Twodimensional simulated spatial gene expression data with a region of interest in orange. (b) Pointwise observations. Orange points are labeled as belonging to the ROI, blue points are outside the ROI, and gray points are unobserved. (c) Estimated EIG for each spatial location (where each design is a single point). (d) Estimated EIG for each horizontal slice design. (e) Synthetic ROI data. (f) Slices chosen after *T* = 5 iterations of running our model. (g) F1 score of predictions after each iteration.

We applied our method to this dataset using the spherical border model (Equation 7). The parameters of interest in this model are the center and radius of the sphere; our approach thus maximizes the EIG in the posterior over these parameters.

For demonstration, we first used a design space where each design was a single point rather than a cross section. We randomly selected 100 points as observations and ran the *EIG* model for one iteration. We then visualized the EIG for each candidate design (each of which was a point in this case). We observed that the EIG was highest for points at the border of the ROI (Figure 3c). This observation implies that, in order to learn the center and radius of the sphere, it is most informative to sample points near the estimated border.

We then extended this experiment to a design space where each design was a line. We ran the *EIG* and *Serial* approaches for *T* = 5 iterations. After each iteration, we visualized the chosen slices, computed predicted labels (ROI or not-ROI) for each point, and computed the predictive performance using the F1 score. We found that the *EIG* approach obtained its maximum predictive performance in fewer iterations compared to the *Serial* approach. This result highlights the usefulness of our design approach when the goal is to localize a region of interest.

### 4.3 Application to Visium data

Next, we applied our experimental design approach to a series of spatial gene expression datasets. We first leveraged spatial transcriptomics data from the 10x Genomics Visium platform (10x Genomics, 2020). This dataset consists of a two-dimensional section from the sagittal-posterior region of a mouse brain. Since a full three-dimensional profile of the brain was not available for this dataset, we considered one-dimensional slices through this two-dimensional tissue as a proof of concept. The goal in this experiment was to characterize the gene expression patterns across the tissue as thoroughly as possible; in other words, the goal was to build an atlas. Thus, we modeled the data with the atlas-building GP regression model (Equation 3), where the parameter of interest is the entire function *f* governing the spatial organization of gene expression.

We ran the *EIG, Serial*, and *Random* design approaches for *T* =10 iterations and visualized the resulting slices. We also sought to quantify the downstream utility of these chosen slices. To do so, we evaluated our ability to impute the gene expression levels at unobserved locations after collecting each slice. We used a GP with an RBF covariance function to make predictions.

We found that the EIG-maximizing slices tended to be the ones that covered the most surface area of the tissue and intersected the most spots, and these slices thoroughly covered the domain of the tissue. (Figure 4b). Moreover, we found that our approach achieved a higher imputation performance across experimental iterations compared to the competing approaches (Figure 4c). This result suggests that our design procedure could be useful for selecting tissue cross sections that will ultimately be used to construct an atlas of the entire tissue.

**Figure 4:**
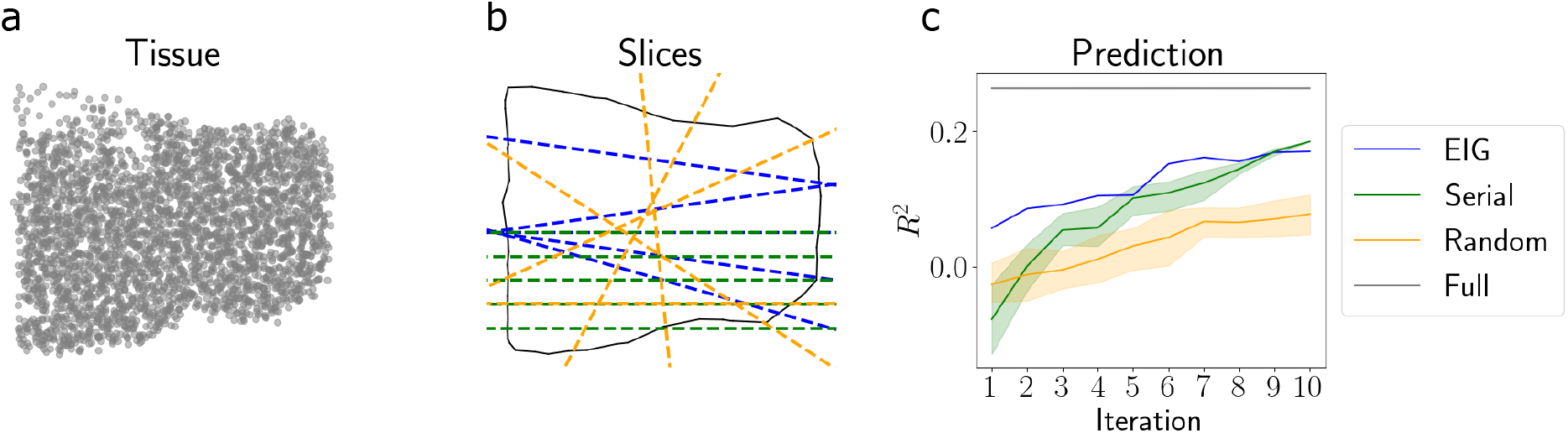
Application to Visium data. (a) Spatial locations of tissue. (b) Slices chosen by each approach after *T* = 5 iterations. The outline of the tissue is shown by the solid black line, and the slices chosen by each approach are shown by the dashed lines. The color legend is in panel (c). (c) Predictive *R*^2^ of the held-out gene expression for both approaches across iterations.

### 4.4 Reconstructing the Allen Brain Atlas

We next applied our experimental design method to three-dimensional spatial gene expression data from the mouse brain in the Allen Brain Atlas (Lein et al., 2007). This dataset contains the expression levels of approximately 20,000 genes in the adult mouse brain and were collected using *in situ* hybridization (ISH). The data are collected as images where the pixel intensity encodes the level of gene expression at each spatial location. The data were collected sagittal sections of the mouse brain that were 200*μm* apart from one another.

While these data were collected with serial slices of the tissue, we sought to answer whether an atlas of equal precision could be constructed with fewer slices using our design approach. To do this, we applied our slicing algorithm to the data and evaluated our ability to impute the gene expression levels of the full atlas. After each slice, we predict the gene expression levels at all unobserved locations and compute the prediction error. For comparison, we compared against two competing slicing strategies: one that takes serial sagittal slices and another that takes random slices. See Appendix A.4.1 for details.

We found that we could reconstruct the atlas within reasonable accuracy with fewer samples than were taken in the original atlas (Figure 5).

**Figure 5:**
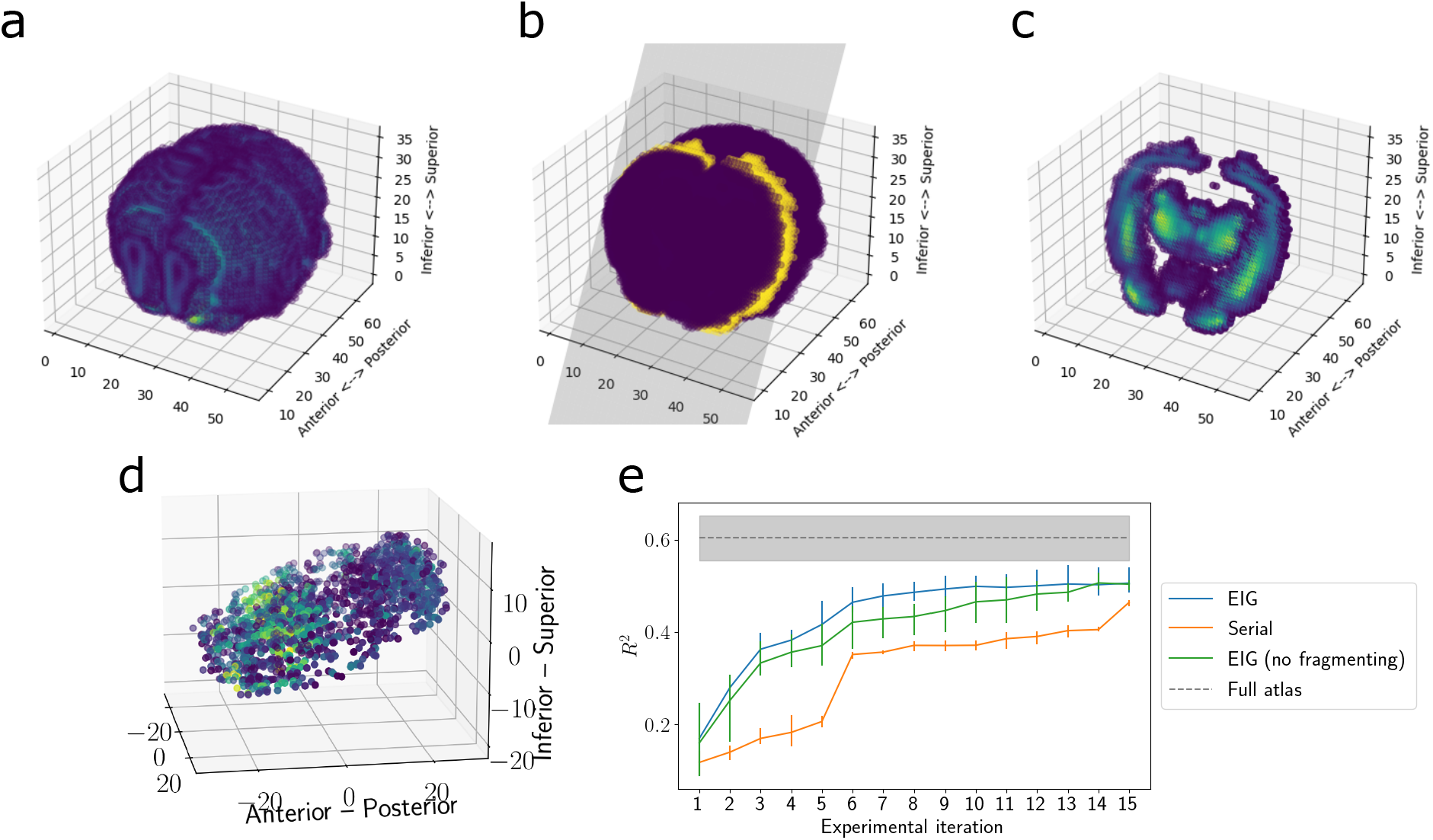
Reconstructing the Allen Brain Atlas. (a) Allen Brain Atlas coordinates colored by the expression of *PCP4*. (b) An example slice through the coordinates. (c) The resulting observations after taking this slice. (d) The slices and observations chosen by the EIG approach. (e) Imputation performance across experimental iterations.

### 4.5 Localizing invasive carcinoma in prostate tissue

As a final application of our experimental design approach, we considered the problem of localizing a tumor within a tissue by taking sequential slices of the tissue. We leveraged another spatial transcriptomics dataset from the 10x Visium platform that profiled a human prostate cancer sample (10x Genomics, 2020). The dataset contains a cross section of the tissue with spatially-resolved transcriptomic data at each location, as well as pathologist annotations of the cancerous tissue region (Figure 6a). We modeled the data with the spherical border model (Equation 7), where again we are interested in estimating the center and radius of the tumor.

**Figure 6:**
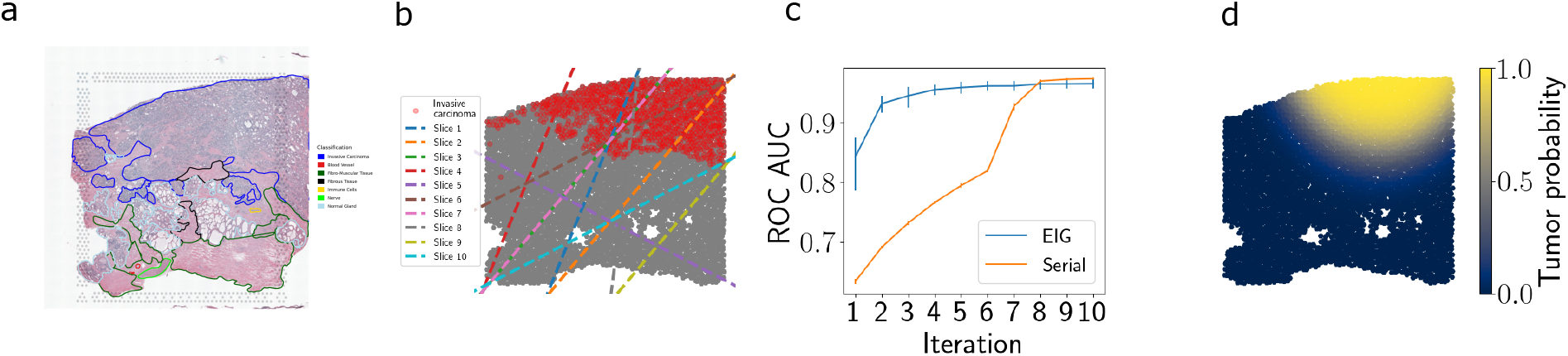
Localizing invasive carcinoma in prostate tissue. (a) Histology image of the tissue section with pathologist annotations overlaid. Image from 10x Genomics website. (b) Slices chosen by the *EIG* method. Cancerous spots are shown in red. (c) F1 score of tumor/healthy label predictions after each iteration of experimental design. (d) Tumor/healthy predictions following five iterations of design. Stronger yellow color indicates spots with higher predicted probability of containing tumorous tissue.

We ran the *EIG* design approach forward for *T* = 5 iterations. On each iteration, we computed model predictions for whether each spatial location corresponded to tumorous or healthy tissue, and we used these predictions to compute the F1 classification score. We found that the classification performance increased rapidly as more slices were collected. This experiment demonstrates the versatility of our design approach to extend to a setting in which a targeted region of the tissue is being localized.

## 5 Discussion

In this paper, we formalized the problem of choosing slice locations for spatial genomics experiments, and we proposed a set of methods for performing experimental design in this setting. We focused on optimizing two study types in particular: constructing an atlas of an entire tissue, and localizing a particular region of a tissue. We applied our method to a range of synthetic datasets, a spatial gene expression dataset from the mouse cortex, and an ISH dataset from the Allen Brain Atlas. In each of these cases, we demonstrated the value of optimizing the locations of cross sections. As spatial genomic profiling technologies evolve, we envision our approach being useful for planning data collection to be as efficient as possible.

### 5.1 Limitations

Our work has several limitations that need to be solved before directly applying our method to design spatial genomics experiments. First, the tools used to obtain slices of tissues may not be precise enough to obtain an exact slice location that is prescribed by our approach.

Second, our approach assumes that a model of the tissue’s shape is available. While such a model is available for many tissue types, it may not be available for less well-studied tissue types. However, even a rough model of the tissue shape suffices for most applications of our model.

### 5.2 Future directions

This work motivates several future directions. First, the experimental design problem could be extended to account for the different types of spatially-resolved measurement technologies available to an experimentalist on any given iteration, as well as their associated costs, spatial resolutions, and levels of precision. Second, there is an opportunity to further formalize the problem of designing experiments with highly structured design constraints, as well as propose new algorithms for optimizing the associated objective. In the current study, we took a simple, brute-force approach to searching over the candidate cross sections, but more efficient search and optimization strategies could be explored.

## Acknowledgements

We thank Addie Minerva, Cate Peña, Bianca Dumitrascu, and Geoffrey Roeder for helpful conversations. AJ and BEE were funded by Helmsley Trust grant AWD1006624, NIH NCI 5U2CCA233195, and NSF CAREER AWD1005627. DC was supported in part by a Google Ph.D. Fellowship in Machine Learning. DL was supported in part by the National Institute of Environmental Health Sciences of the National Institutes of Health under award number P30ES010126 University of North Carolina Center for Environmental Health and Susceptibility. The content is solely the responsibility of the authors and does not necessarily represent the official views of the National Institutes of Health.

## A Appendix

### A.1 Supplementary methods

#### A.1.1 Alternate form of IG

Consider a setting in which we are choosing a design *x*. Suppose our model for the data *y* is parameterized by *θ*, which we wish to perform inference on. The information gain after choosing design *x* is given by the reduction in differential entropy from prior to posterior:

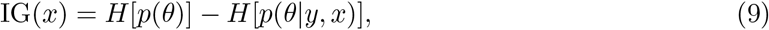

where 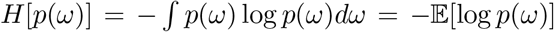 is the entropy of a random variable *ω*. Expanding the posterior in Equation 9 using Bayes rule, we have

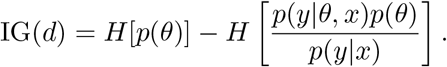

Note that

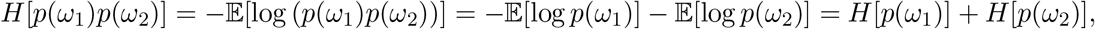

allowing us to split up the second term:

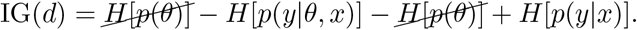

This yields the alternative form of IG due to the symmetry of mutual information (MI):

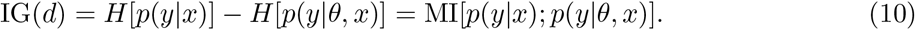

This is easily extended to the setting where we’re choosing the design *x* for the *t*th experimental iteration, and we have already observed data **Y**_1:*t*-1_. In this case, our “prior” is the posterior for *θ* given **Y**_1:*t*-1_, and the IG calculation is nearly identical. This yields

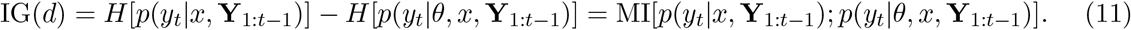

#### A.1.2 EIG for GP regression

Consider the GP regression model:

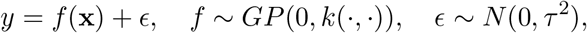

where 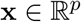. Suppose our design space is 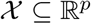, and on each iteration we choose a single design **x**.

Recall the differential entropy of a *D*-dimensional multivariate normal distribution with mean vector **m** and covariance matrix **C**:

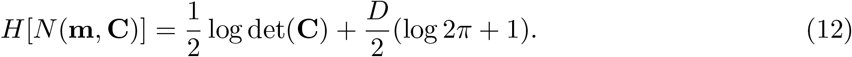

We can plug the entropy into the equation for the EIG (Equation 10) to compute the EIG under the GP regression model.

On the first experimental iteration (before observing any data), we can compute the EIG using Equation 10 as follows:

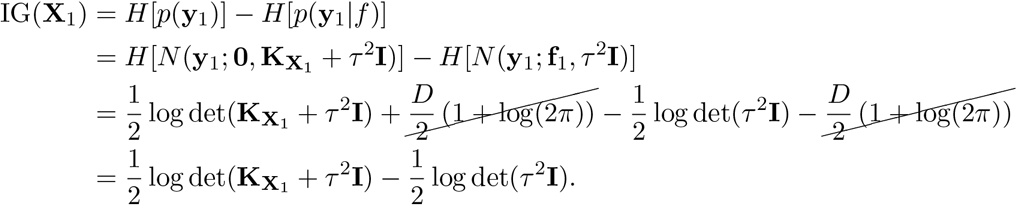

Factoring out *τ*^2^**I** in the log of the first term leads us to cancel the second term as follows:

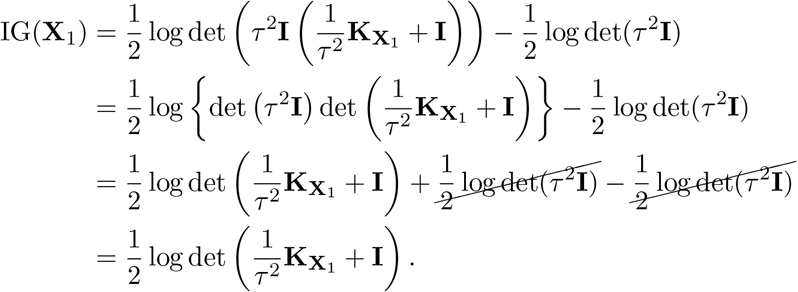

On experimental iteration *t* > 1, we can compute the IG using Equation 11:

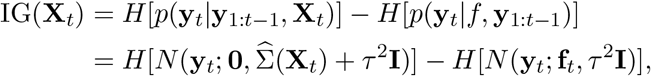

where 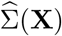 is the posterior covariance of the GP, given by

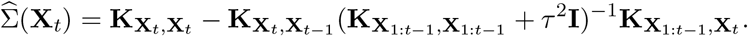

Following a similar series of calculations, we obtain

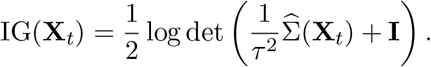

#### A.1.3 Variational inference approach

In the border-finding application, we approximate the posterior *p*(*θ*|**y**) using variational inference (Hoffman et al., 2013), where the variational family is specified as the set of multivariate Gaussians with diagonal covariance matrices. Specifically, *q*(*θ*) = *N_K_*(*θ*|***η***,diag(***ψ***)), where *K* is the number of model parameters, ***η*** is the variational mean, and **ψ** is the vector of variational marginal variances. We then minimize the KL divergence between the approximate posterior distribution and the posterior distribution with respect to the variational parameters, *φ* = **{*η*, *ψ*}**. This is equivalent to maximizing the evidence lower bound (ELBO). To see, this note that the KL divergence from *q*(*θ*) to *p*(*θ*|**y**) can be split into the log evidence and the ELBO:

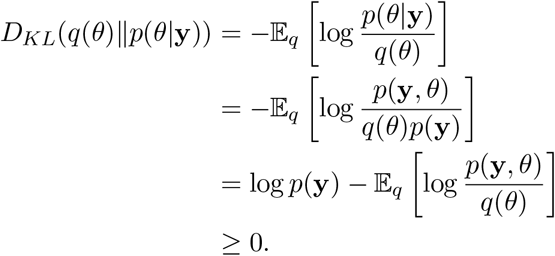

The KL divergence is nonnegative, so we obtain a lower bound on the log evidence:

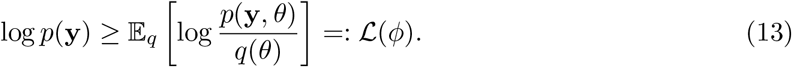

We maximize the ELBO 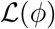 with respect to *φ* using stochastic variational inference (Hoffman et al., 2013).

### A.2 Data availability

#### Synthetic data

Code for generating synthetic datasets can be found in the GitHub repository: https://github.com/andrewcharlesjones/spatial-experimental-design.

#### Visium mouse brain data

Visium data were obtained from the 10x Genomics website. Data for the one slice was downloaded from the “Datasets” page. Specifically, spatial gene expression was downloaded from the page called *Mouse Brain Serial Section 1 (Sagittal-Posterior)*.

#### Allen Brain Atlas data

The *in situ* hybridization data from the Allen Brain Atlas was downloaded using the brainrender Python API (Claudi et al., 2021). The GitHub repository for the API is available at https://github.com/brainglobe/brainrender. We used data from the study with experiment ID 79912613.

#### Visium prostate cancer data

Visium data were obtained from the 10x Genomics website from the Datasets page titled *Human Prostate Cancer, Adenocarcinoma with Invasive Carcinoma (FFPE)*.

### A.3 Code availability

Code for all data preprocessing, synthetic data generation, and experiments can be found in the GitHub repository.

### A.4 Experimental details

#### A.4.1 Allen Brain Atlas experiment

##### Data preprocessing

To filter out voxels outside of the tissue, we removed spatial locations with intensity less than two. The spatial coordinates were centered to have a mean of zero, and the intensities for each gene were standardized by subtracting the mean and dividing by the standard deviation.

#### A.4.2 Visium prostate cancer experiment

The provided graph-based cluster labels were used to label the spots containing carcinoma cells (spots belonging to Cluster 1 were labeled as carcinoma).

### A.5 Supplementary figures

**Supplementary Figure 1:**
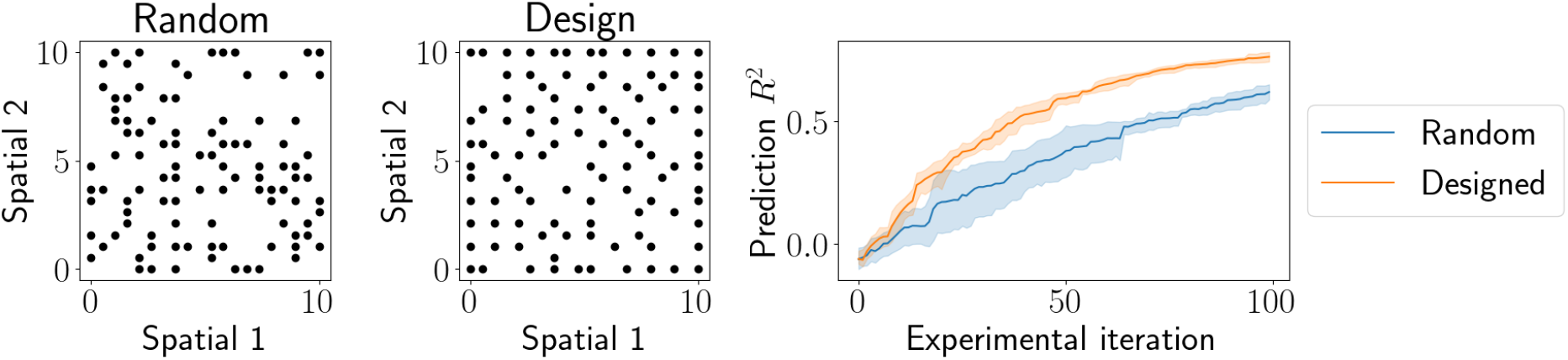
Experimental design with one-dimensional design space. (a) Designs after *T* = 100 iterations for the *Random* approach. (b) Designs after *T* = 100 iterations for the *EIG* approach. (c) Imputation performance after each new observation.

## References

10x Genomics (2020). Mouse Brain Serial Sections (Sagittal-Posterior), Spatial Gene Expression Dataset by Space Ranger 1.1.0, 10x Genomics, (2020, June 23).

Baker, E. A. G., Schapiro, D., Dumitrascu, B., Vickovic, S., and Regev, A. (2022). Power analysis for spatial omics. bioRxiv.

Box, J. F. (1980). RA Fisher and the design of experiments, 1922-1926. The American Statistician, 34(1):1–7.

Buzug, T. M. (2011). Computed tomography. In Springer handbook of medical technology, pages 311–342. Springer.

Camerlenghi, F., Dumitrascu, B., Ferrari, F., Engelhardt, B. E., and Favaro, S. (2020). Nonparametric Bayesian multiarmed bandits for single-cell experiment design. The Annals of Applied Statistics, 14(4):2003–2019.

Chaloner, K. and Verdinelli, I. (1995). Bayesian experimental design: A review. Statistical Science, pages 273–304.

Claudi, F., Tyson, A. L., Petrucco, L., Margrie, T. W., Portugues, R., and Branco, T. (2021). Visualizing anatomically registered data with brainrender. Elife, 10:e65751.

Fisher, R. (1926). The arrangement of field experiments. Journal of the Ministry of Agriculture of Great Britain, 33(48):503–513.

Foster, A., Ivanova, D. R., Malik, I., and Rainforth, T. (2021). Deep adaptive design: Amortizing sequential Bayesian experimental design. In International Conference on Machine Learning, pages 3384–3395. PMLR.

Foster, A., Jankowiak, M., Bingham, E., Horsfall, P., Teh, Y. W., Rainforth, T., and Goodman, N. (2019). Variational Bayesian optimal experimental design. Advances in Neural Information Processing Systems, 32.

Foster, A., Jankowiak, M., O’Meara, M., Teh, Y. W., and Rainforth, T. (2020). A unified stochastic gradient approach to designing Bayesian-optimal experiments. In International Conference on Artificial Intelligence and Statistics, pages 2959–2969. PMLR.

Hoffman, M. D., Blei, D. M., Wang, C., and Paisley, J. (2013). Stochastic variational inference. Journal of Machine Learning Research.

Kierszenbaum, A. L. and Tres, L. (2015). Histology and Cell Biology: an introduction to pathology. Elsevier Health Sciences.

Lein, E. S., Hawrylycz, M. J., Ao, N., Ayres, M., Bensinger, A., Bernard, A., Boe, A. F., Boguski, M. S., Brockway, K. S., Byrnes, E. J., et al. (2007). Genome-wide atlas of gene expression in the adult mouse brain. Nature, 445(7124):168–176.

Lindley, D. V. (1956). On a measure of the information provided by an experiment. The Annals of Mathematical Statistics, 27(4):986–1005.

Mantri, M., Scuderi, G. J., Abedini-Nassab, R., Wang, M. F., McKellar, D., Shi, H., Grodner, B., Butcher, J. T., and De Vlaminck, I. (2021). Spatiotemporal single-cell RNA sequencing of developing chicken hearts identifies interplay between cellular differentiation and morphogenesis. Nature Communications, 12(1):1–13.

Masoero, L., Camerlenghi, F., Favaro, S., and Broderick, T. (2022). More for less: predicting and maximizing genomic variant discovery via Bayesian nonparametrics. Biometrika, 109(1):17–32.

Moses, L. and Pachter, L. (2022). Museum of spatial transcriptomics. Nature Methods, 19(5):534–546.

Quake, S. R. (2022). A decade of molecular cell atlases. Trends in Genetics.

Rainforth, T., Cornish, R., Yang, H., Warrington, A., and Wood, F. (2018). On nesting Monte Carlo estimators. In International Conference on Machine Learning, pages 4267–4276. PMLR.

Rozenblatt-Rosen, O., Stubbington, M. J., Regev, A., and Teichmann, S. A. (2017). The Human Cell Atlas: from vision to reality. Nature, 550(7677):451–453.

Russell, E. (1926). Field experiments: How they are made and what they are. Journal of the Ministry of Agriculture, 32(989):1001.

Saviano, A., Henderson, N. C., and Baumert, T. F. (2020). Single-cell genomics and spatial transcriptomics: Discovery of novel cell states and cellular interactions in liver physiology and disease biology. Journal of Hepatology, 73(5):1219–1230.

Schmid, K. T., Cruceanu, C., Böttcher, A., Lickert, H., Binder, E. B., Theis, F. J., and Heinig, M. (2020). Design and power analysis for multi-sample single cell genomics experiments. bioRxiv.

Shekhar, K. and Sanes, J. R. (2021). Generating and using transcriptomically based retinal cell atlases. Annual Review of Vision Science, 7:43–72.

Smith, E. A. and Hodges, H. C. (2019). The spatial and genomic hierarchy of tumor ecosystems revealed by single-cell technologies. Trends in Cancer, 5(7):411–425.

Steinberg, D. M. and Hunter, W. G. (1984). Experimental design: review and comment. Technometrics, 26(2):71–97.

Sunkin, S. M., Ng, L., Lau, C., Dolbeare, T., Gilbert, T. L., Thompson, C. L., Hawrylycz, M., and Dang, C. (2012). Allen Brain Atlas: an integrated spatio-temporal portal for exploring the central nervous system. Nucleic Acids Research, 41(D1):D996–D1008.

Svensson, V., Natarajan, K. N., Ly, L.-H., Miragaia, R. J., Labalette, C., Macaulay, I. C., Cvejic, A., and Teichmann, S. A. (2017). Power analysis of single-cell RNA-sequencing experiments. Nature Methods, 14(4):381–387.

Yoosuf, N., Navarro, J. F., Salmén, F., Ståhl, P. L., and Daub, C. O. (2020). Identification and transfer of spatial transcriptomics signatures for cancer diagnosis. Breast Cancer Research, 22(1):1–10.

Zhang, M., Eichhorn, S. W., Zingg, B., Yao, Z., Cotter, K., Zeng, H., Dong, H., and Zhuang, X. (2021). Spatially resolved cell atlas of the mouse primary motor cortex by merfish. Nature, 598(7879):137–143.

